# Dynamically driven correlations in elastic net models reveal sequence of events and causality in proteins

**DOI:** 10.1101/2024.01.15.575718

**Authors:** Albert Erkip, Burak Erman

## Abstract

Protein dynamics orchestrate allosteric regulation, but elucidating the sequence of events and causal relationships within these intricate processes remains challenging. We introduce the Dynamically Perturbed Gaussian Network Model (DP-GNM), a novel approach that uncovers the directionality of information flow within proteins. DP-GNM leverages time-dependent correlations to achieve two goals: identifying driver and driven residues and revealing communities of residues exhibiting synchronized dynamics. Applied to wild type and mutated structures of Cyclophilin A, DP-GNM unveils a hierarchical network of information flow, where key residues initiate conformational changes that propagate through the protein in a directed manner. This directional causality illuminates the intricate relationship between protein dynamics and allosteric regulation, providing valuable insights into protein function and potential avenues for drug design. Furthermore, DP-GNM’s potential to elucidate dynamics under periodic perturbations like the circadian rhythm suggests its broad applicability in understanding complex biological processes governed by environmental cycles.

## Introduction

Proteins are dynamic and complex molecular systems that carry out their diverse functions through orchestrated thermal fluctuations. These fluctuations are not random but rather organized and consequential, enabling proteins to adopt conformational flexibility and engage in intricate functional interactions with other molecules. Proteins constantly experience perturbations, both intrinsic and extrinsic, arising from both stochastic and deterministic sources. Intrinsic perturbations include mutations and post-translational modifications, while extrinsic perturbations stem from ligand or protein binding events. Both types of perturbations modulate the protein’s energy landscape, resulting in alterations to its dynamics and equilibrium properties. Static effects of perturbations in proteins are widely studied ^1, 2^. The present work is on the dynamic allosteric effects of perturbations in proteins, based on the Langevin equation solution of the dynamics of a Gaussian Network Model (GNM)^3^. The dynamic version of the GNM that we use here was introduced^4^ in 2004, where a single-frequency cosine perturbation was applied to a specific mode of the GNM and the response showed significant synchronicity between different modes. In 2021, the modal cosine perturbation approach was extended^5^ to perturbation of single residues, allowing one to probe the dynamic response of the entire protein upon perturbing a specific residue. This mimics perturbation-response dynamics observed in allosteric regulation. The present paper introduces the ‘Dynamically Perturbed GNM Model’ (DP-GNM), which builds upon the two previous models where we now focus on the causal relationships introduced into the dynamics. Causality, defined by the difference in influence between residue i on j and j on i over time, is a critical aspect of protein dynamics as it forms the foundation for information transfer and allostery. Recently, Scaramozzino et al.^6, 7^ extended the Langevin equation solution to another elastic network model, the Anisotropic Network Model (ANM), and analyzed protein fluctuations in response to random external forces, validating our previous results for the GNM.

An essential aspect of causality is the sequence of events or the directionality of allostery. A diverse array of methods has emerged to determine the key residues involved in allosteric modulation^8-17^, only few computational papers elaborated on the sequence of events and causality in this process^18-22^. Interestingly, recent experimental work^23-26^ clearly shows the importance of causality in dynamic allostery.

We show that by externally perturbing a protein, and calculating time correlation function of residue fluctuations, we can identify the residues that exert a stronger influence on the response of other residues, i.e. we can determine the direction of information flow. An extreme analogy of directional information flow would be the electric switch and the lightbulb where on/off information flows from the switch to the lightbulb but no information flows from the lightbulb to the switch. Direction of information flow cannot be determined with the use of the static picture of proteins. The ‘driver-driven’ terminology aptly captures the directional nature of information flow in proteins, with drivers acting as switches and driven residues receiving the signals.

The DP-GNM, which we introduce in this study focusses on phase differences between fluctuations of residues, hence it becomes possible to answer the question ‘Which residues move in phase with the fluctuations of the perturbed residue?’ The answer to this question helps in the clear identification of communities and networks of allosteric residues in the protein.

Crucially, the directionality identified by DP-GNM is not an inherent property of the unperturbed protein, but rather emerges as a consequence of the applied perturbation and its interaction with the protein’s dynamic landscape. Moreover, this directionality exhibits a periodic reversal between successive time intervals of (*n* −1)*π* < *ωτ* < *nπ, n* = 1, 2, 3,… where *ωτ* represents the product of applied frequency *ω* and delay time*τ*. This periodicity highlights the dynamic nature of information flow within proteins, where the roles of driver and driven residues can shift depending on the frequency of perturbation and timescale of observation.

To illustrate how one can use time-dependent perturbations to study allostery and the sequence of events in proteins, we examine the system Cyclophilin A (CypA) which is an important allosteric protein that catalyzes the conversion of cys-trans transition of proline residues. We study the wild type cyclophilin A and two mutated structures.

## Theory

In this section, we introduce the Dynamically Perturbed Gaussian Network Model, DP-GNM, and Causal relations. The details of the mathematical derivations are presented in the Supplementary material section. Here we give only the results of the detailed derivations.

### 1. Formulation of the problem

The time correlation *C*_*ij*_(*τ*) of the fluctuations Δ*R* of two residues i and j is defined as

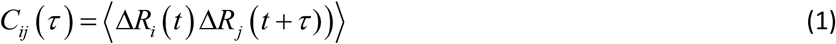

where *τ* is the time delay, i and j denote the residue indices and angular brackets denote an average. The fluctuations are vectorial quantities and the product represents the scalar product. GNM assumes that fluctuations are spherically symmetric, therefore considering only one component is sufficient. *C*_*ij*_(*τ*) represents the effect of the fluctuations of i on subsequent fluctuations of j. If *C*_*ij*_(*τ*) is larger than *C*_*ji*_(*τ*), for example, then one can assign directionality to the system and say that fluctuations of i drive the fluctuations of j. This is how the DP-GNM introduces causality into allostery.

The time evolution of the protein of n residues obeys the Langevin equation

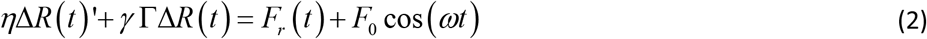

where, Δ*R*(*t*), the instantaneous fluctuation vector, is an nx1 vector with the ith entry referring to the instantaneous fluctuation of the ith residue. For anisotropic fluctuations, each entry of Δ*R*(*t*) will have three components in three coordinate directions. Here, we consider a single value, which may be assumed as the average of the three since we adopt the isotropic representation^3^ of the GNM for simplicity. The prime in Eq. 2 shows differentiation with respect to time, *η* is the friction coefficient, *γ* is a characteristic spring constant and Γ is the nxn dimensionless Kirchoff matrix where the ij’th term is -1 if residues i and j are within a cutoff distance of each other and interacting, or zero otherwise. Each diagonal term equates to the negative sum of the corresponding row. In the theory, the ratio 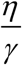will appear, which we call the effective viscosity, and denote by *ζ* .The right hand side represents the sum of a random force *F*_*r*_(*t*) and a deterministic force *F*_0_ cos(*ωt*) on the protein. In our previous work, we showed^4^ that the solution of the Langevin equation where the random force is white noise that obeys ⟨*F*_*i*_(*t*) *F*_*j*_(*t*’)⟩ = *ηk*_*B*_*Tδ*_*ij*_*δ*(*t* − *t*’) leads to the GNM model. Here, *kB* is the Boltzmann constant, *T* is the temperature, *δ*_*ij*_ is the Kronecker delta which is equal to 1 if i and j are equal and zero otherwise, and *δ*(*t* − *t*’) is the Dirac function which is zero if the argument is nonzero and infinity if the argument is zero. In the present work we superpose on the random force a deterministic component, *F*(*t*) = *F*_0_ cos*ωt*, where *ω* is the frequency of the applied force and *F*_0_ is a nx1 vector whose ith entry is the force on the ith residue. In this study, we perturb the GNM model by simultaneously applying a white noise generated random force and a time-varying force with a cosine profile. The solution of Eq. 2 is obtained as the superposition of three functions, Δ*R*(*t*) = Δ*R*_*r*_(*t*) + Δ*R*_*p*_(*t*) + *Ce*(−*ζ t*Γ) where Δ*R*_*r*_(*t*) is the solution for the random force and Δ*R*_*p*_(*t*) is the solution for the deterministic force. The solution to the random force is given in Reference ^4^ which is also presented in the Supplementary Material. The solution to the deterministic force is

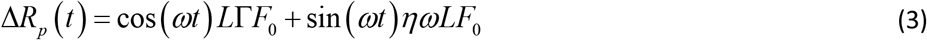

where *L* is an nxn matrix given by

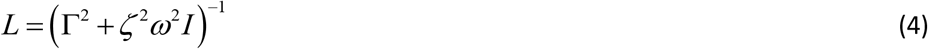

where, the ij’th component is

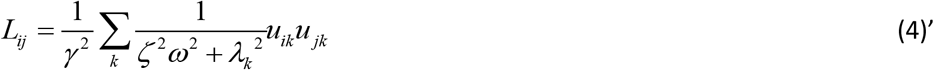

with *ζ* = *η*/*γ*, i.e., the effective vicosity, and *λ*_*k*_ and *u*_*ik*_ showing the kth eigenvalue and the i’th entry of the kth eigenvector of the Γ matrix, respectively. We are interested not in the fluctuations Δ*R*_*r*_(*t*) and Δ*R*_*p*_(*t*) in the solution of the Langevin equation, but in the time correlation function obtained from them:

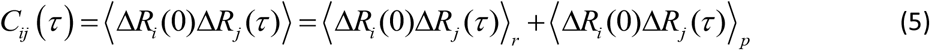

The subscripts r and p refer to Δ*R*_*i*_(*t*)_*r*_ and Δ*R*_*i*_(*t*)_*p*_, respectively. The averages are performed on the assumption that the system is ergodic and time averages are equal to ensemble averages. In Eq. 5, the cross terms ⟨Δ*R*_*i*_ (0)_*r*_ Δ*R*_*j*_(*τ*)_*p*_⟩, ⟨Δ*R*_*i*_(0)_*p*_ Δ*R*_*j*_(*τ*)*R*⟩ are zero since the average of a zero mean random and a deterministic continuous function is zero. Moreover, the exponential term *Ce*^(−*ζt*Γ)^ does not contribute as it decays exponentially as *t* →∞. We give the derivation of the explicit forms of these functions in the Supplementary Material Section. Here, we show only the results:

i. Time correlation for the unperturbed protein: The expression for time delayed pair correlations in an unperturbed GNM is given^4, 27^

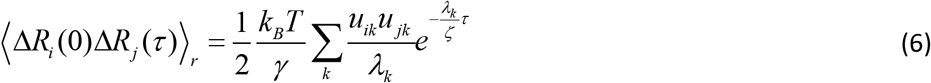

This correlation function is symmetric in i and j.
ii. Time correlation in perturbed network: When residue p is perturbed, the correlation between i at time 0 and j at time *τ* is given as (See Supplementary material section for derivation)

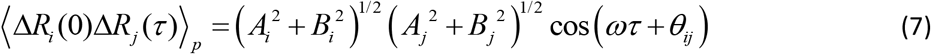

where,

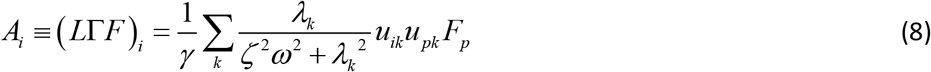

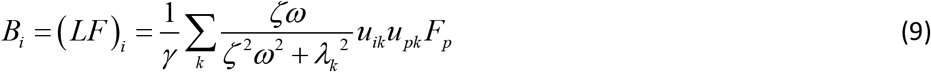

and

*θ*_*ij*_ = *θ*_*j*_ −*θ*_*i*_ represents the phase shift with *θ*_*i*_ = arctan 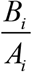, and related to *A*_*i*_ and *B*_*i*_ with

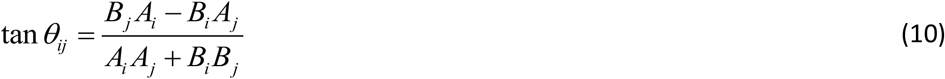

Since *θ*_*ij*_ is antisymmetric, ⟨Δ*R*_*i*_(0)Δ*R*_*j*_(*τ*)⟩_*p*_ will not be equal to ⟨Δ*R*_*j*_(0)Δ*R*_*i*_(*τ*)⟩_*p*_. When p is perturbed, response of i will contain a phase difference and response of j will have another phase difference. Therefore correlation of i at time zero and of j at time *τ* will contain a phase difference *θ*_*ij*_. However, if j is observed first and i after a time*τ* the phase difference will be −*θ*_*ij*_. It is this difference that leads to causality. In the calculations, the perturbed residue p may be the residue i or j.

### 2. Dimensional analysis and choice of system variables

The theoretical model is a coarse-grained model and the correspondence of its parameters to the real system should be considered. Denoting the dimensions of time, length and force by *T, L* and *F*, respectively, the dimensions of the friction constant *η*, the spring constant *γ* and the applied frequency *ω* respectively are *FT*/*L, F*/*L* and 1/ *T*. The effective viscosity *ζ* will then have the dimension of time and the eigenvalues will be dimensionless. Equations 7-10 which we use will all be expressed in terms of three dimensionless parameters, *ωτ, ζω* and the set of eigenvalues *λ*_*i*_. The average of the eigenvalues 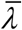 is related to the sum of the diagonal elements of Γ by the relation

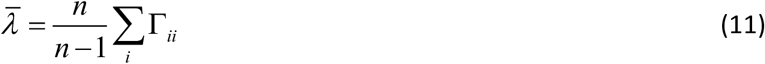

As an example, taking a cutoff value 7.0 Å for Cyclophilin A in Eq. 11 gives an average number of 8.2 for both sides. If dynamics close to the unperturbed state is of interest, then values of *ζ* ^2^*ω* ^2^ should be taken such that 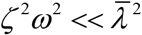. The average of the smallest 10 eigenvalues of Γ, which shows large scale collective motions, is approximately 1.0. Therefore a value of *ζ* ^2^*ω*^2^ ≤ 1 is representative of large scale, slow dynamics.

Relaxation of correlations in the random force case may be written in terms of the dimensionless parameters as

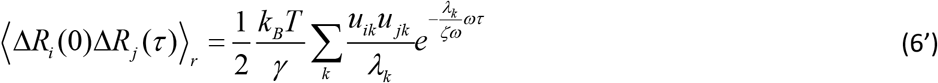

Equation 6’ may be used as a benchmark for comparing the dynamics of the perturbed and the unperturbed protein.

In the formulation, *γ* has dimensions of *FL*^−1^. In order to make the model easily scalable and to remove the dimensionality, we equate *γ* to unity in all the calculations below.

### 3. Factors affecting the causality of information transfer between a residue pair

The time correlation given by eq. 7 is a cosine function, which is shifted with respect to the applied force. The amount of shift depends on the parameters of the system. The numerator on the right hand side which induces the shift is antisymmetric in i and j and depends on the effective friction coefficient, frequency, the eigenvectors of the Γ matrix and the applied force. Due to the presence of the antisymmetric *θ*_*ij*_, *C*_*ij*_(*τ*) and *C*_*ji*_(*τ*) are not equal. The inequality *C*_*ij*_(*τ*) ≠ *C*_*ji*_(*τ*)is the source of causality between i and j. If ⟨Δ*R*_*i*_(*t*) Δ*R*_*j*_(*t* +*τ*)⟩ > ⟨Δ*R*_*j*_(*t*) Δ*R*_*i*_(*t* +*τ*)⟩, then the fluctuations of residue i at time t has more influence on the future values of the fluctuations of j at *t* +*τ*. Equation 7 is a periodic function. The front factor on the right hand side does not depend on the order ij or ji, the only dependence being on *θ*_*ij*_. Therefore, examination of the cos(*ωτ* ± *θ*_*ij*_) term in Eq.7 will shed light on directionality of correlations. The two cosine curves will be shifted relative to each other by an amount of 1± cos(*θ*_*ij*_), where the ± sign depends on the sign of *θ*_*ij*_. For positive *θ*_*ij*_, *C*_*ij*_(*τ*) = cos(*ωτ* + *θ*_*ji*_) will be larger than *C*_*ji*_(*τ*) = cos(*ωτ* + *θ*_*ij*_) for the intervals 0 < *ωτ* < *π*, 2*π* < *ωt* < 3*π* … This is sketched in the inset in Figure 1. For illustrative purposes we use the following values for *θ*_*ij*_ = 1 and *θ*_*ji*_ = −1. In the figure, the abscissa shows the values of *ωτ* and the ordinate values for *C*_*ij*_(*τ*) (solid curve) and *C*_*ji*_(*τ*) (dot dashed curve). In the present study, we assume low frequencies such that the system is in the first interval 0 < *ωτ* < *π* as sketched in the main part of Figure 1.

**Figure 1.**
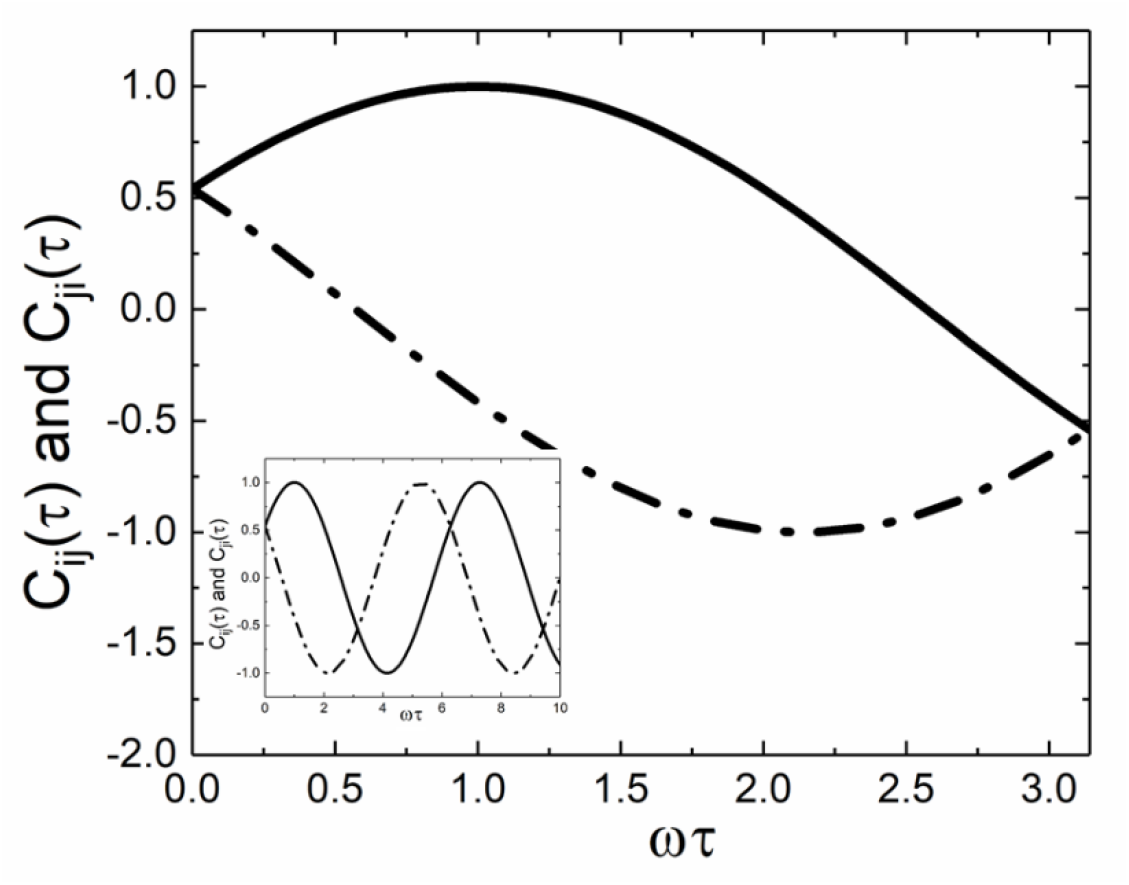
Sketch of cos(*ωτ* + *θ*_*ji*_) (solid curve) and cos(*ωτ* + *θ*_*ij*_) (dot dashed curve) both in the inset and the main figure. The solid curve shows the values of the correlation *C*_*ij*_(*τ*) when residue i is perturbed.

We now consider the low frequency region, i.e. the region 0 < *ωτ* < *π* in Figure 1. Examination of equations 8 and 9 shows that the shift is proportional to the difference *B*_*j*_*A*_*i*_ − *B*_*i*_*A*_*j*_. The sign of this function which determines the direction of information transfer between i and j depends on the eigenvalues and the eigenvectors of the gamma matrix, i.e., the topology of the protein structure.

Finally, we note that the direction of information transfer through correlations reverses periodically as a function of frequency and delay time. Therefore, there is no absolute information transfer direction between two residues. In the present study, we consider slow frequencies and short delay times, i.e., the regime where 0 < *ωτ* < *π*.

### 4. Identifying switch and receptor residues

If *C*_*ij*_(*τ*) − *C*_*ji*_(*τ*) > 0 then residue i acts as the driver of j. A large positive value of the sum

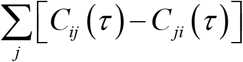 indicates that residue i drives several residues and hence acts as a switch in the protein. Conversely, a large negative value of this sum indicates that the residue i is a driven residue of the protein, analogous to the lightbulb example. Identification of switch and lightbulb residues of the protein gives significant information on the allosteric behavior of the protein. We show the significance and importance of this in the results section.

### 5. Identifying residues that move synchronously; Communities

If two residues i and j move synchronously, then the phase difference between their motions, given by Eq. 10, will vanish. Identifying the set of residues that move synchronously with the perturbed residue may reveal valuable information about protein dynamics and potentially allosteric pathways. These synchronously moving residues form a network directly or indirectly connected to the perturbed residue. Their movements are dynamically correlated, meaning a change in one affects the others, often due to physical interactions or conformational changes propagating through the protein. This network may be involved in protein functions. Residues directly coupled to the perturbation site might be critical for substrate binding, enzyme activity, or other mechanisms. Synchronous movements suggest communication pathways within the protein structure. The network likely transmits information or signals from the perturbation site to other functional regions. If the perturbation site is functionally relevant, its connected network potentially forms the communication route for allosteric effects. Alternatively, we can consider the relationship between synchronously moving parts and communities. Proteins often exhibit a modular architecture, where semi-autonomous sub-units called communities perform specific functions. Identifying synchronously moving residues can tell us which communities are functionally coupled. For example, communities involved in substrate binding and catalysis might move together when the substrate binds. Communities are not static entities; they dynamically interact and communicate to fulfill complex protein functions. By analyzing synchronous movements, we can learn about the communication pathways between communities, crucial for understanding how different functional modules work together. Allosteric regulation relies on long-range signal propagation within the protein. Communities can serve as “communication hubs” within this network. Identifying synchronously moving residues can help us map these allosteric pathways, highlighting the communities mediating signal transmission from the effector site to the functional output. Allosteric effects can result in the reorganization of communities. For example, ligand binding might reconfigure interactions between communities, altering their dynamics and influencing overall protein function. Therefore, studying synchronous movements can reveal these dynamic changes and their functional consequences.

## Application of the DP-GNM to Cyclophilin A

### 1. The structure of Cyclophilin A and the allosteric network residues

In this section we apply the results of the DP-GNM to a widely studied protein Cyclophilin A, a 163 residue protein, which binds its substrate at its active site and catalyzes the cys-trans transition of the proline residue of its substrate. The protein Data Bank code is 3k0n. The residues participating in catalytic activity were determined using nmr relaxation experiments^28^ and studied using molecular dynamics simulations^29, 30^. Simulations showed that catalytic activity of CypA could be explained by changes in motions of residue Phe113 on a ns–μs timescale. The three dimensional structure of CypA is shown in Figure 2.

**Figure 2.**
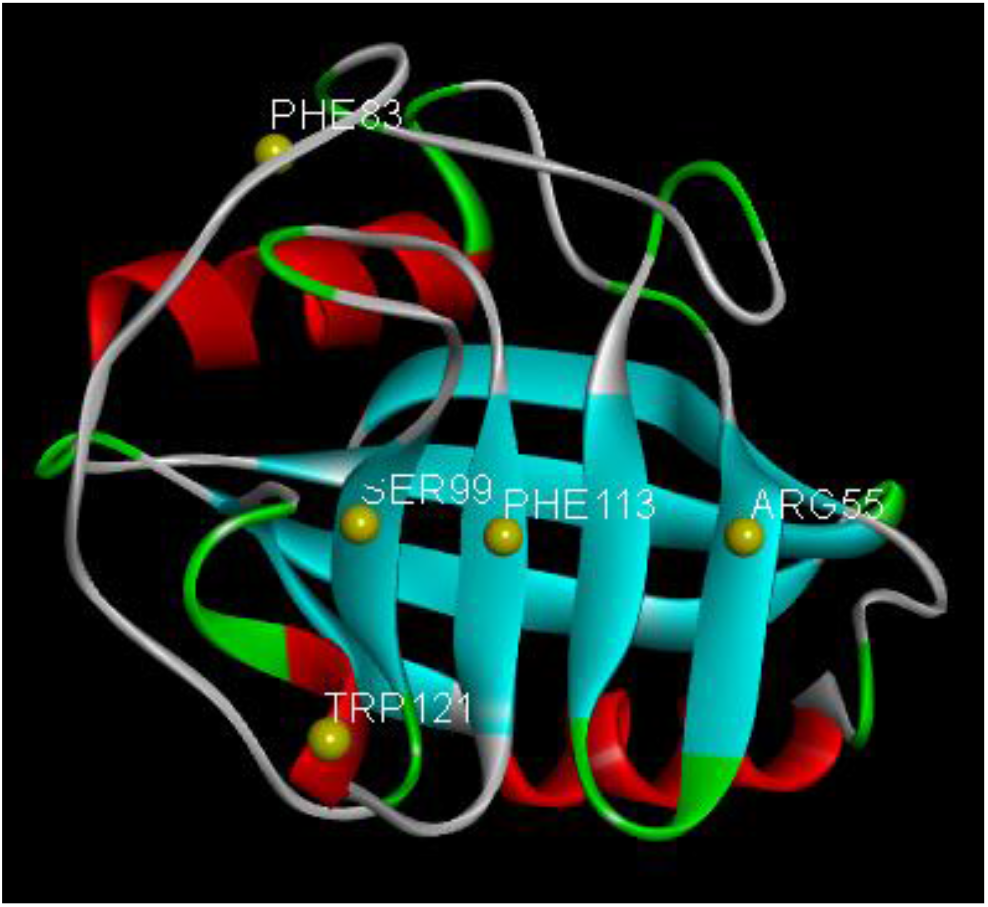
Ribbon diagram of Cyclophilin A, 3K0N.pdb, where yellow spheres show the alpha carbons of the allosteric network residues.

The residues that are known^28, 29^ to constitute the allosteric network, Arg55, Phe83, Ser99, Phe113, Trp121 are identified by their alpha carbons as yellow spheres. Arg55 is the key active site residue, which has the catalytic activity of converting a proline of a substrate between cis and trans states. The allosteric interactions between Arg55 and the remaining allosteric network residues are the key components of the catalytic activity. Although the allosteric activity of Cyclophilin A has been addressed in several papers^9, 29-38^ causality and directional information transfer have not been considered in detail.

### 2. Determining the driver and driven residues in pair interactions

According to Eq. 7 when residue i is perturbed, its correlation with j at a later time will differ from the correlation of j with the perturbed i at a later time. If | *C*_*ij*_(*ωτ*) | > | *C*_*ji*_(*ωτ*) |, we call residue i the driver and j the driven. In order to understand the driver-driven relations between residue Arg55, the major player in catalysis, and the residues of the allosteric network, we perturbed residue Arg55 and calculated the ensuing correlations. In Figure 3, we show the time correlation functions between Arg55 and other members of the allosteric pathway. It is interesting to see that the perturbed residue does not always become the driver as would normally be expected. Calculations with the remaining residues of the protein support this observation.

**Figure 3.**
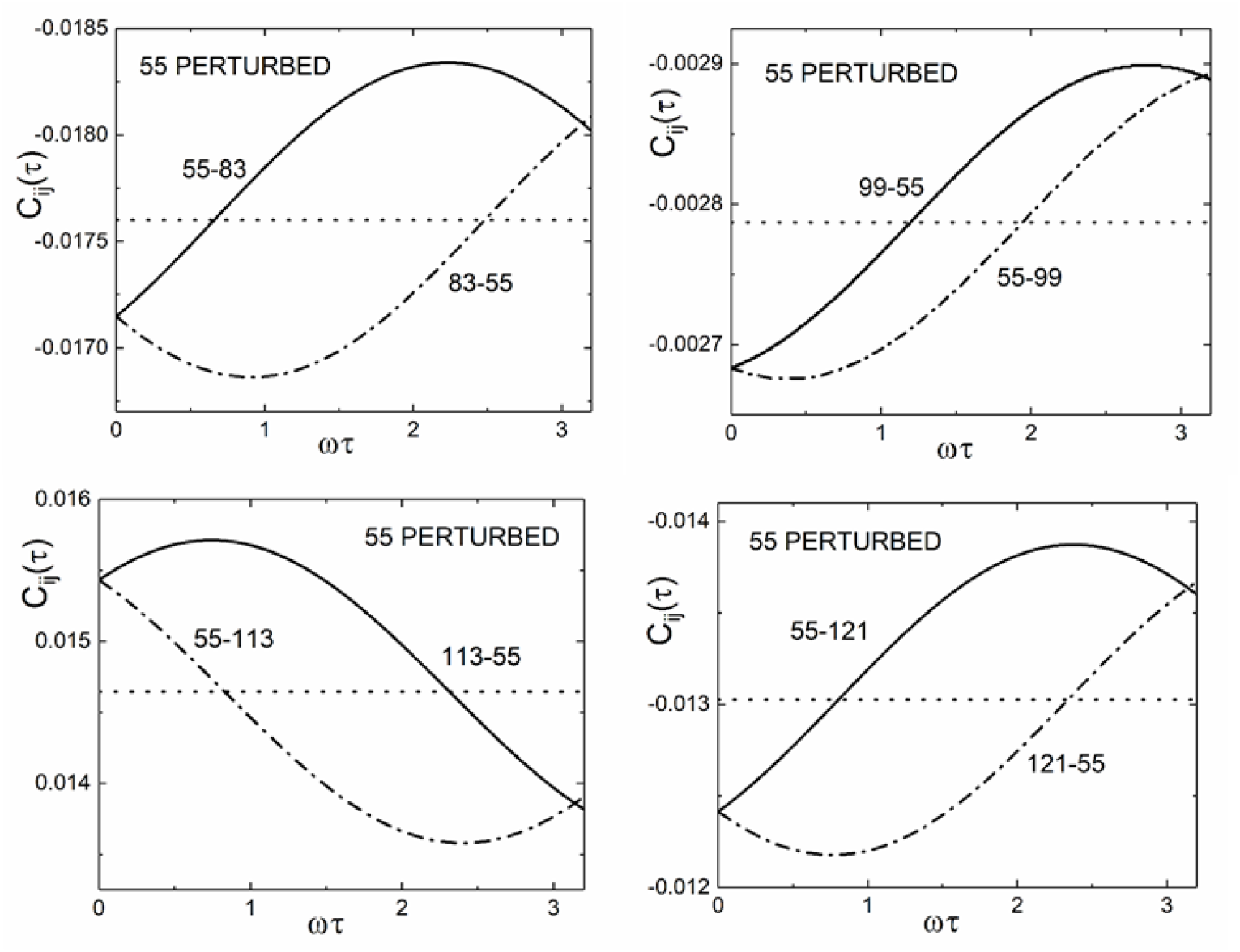
Time correlations between residue Arg55 and residues Phe83, Ser99, Phe113 and Trp121 as function of *ωτ*. In all panels, residue Arg55 is perturbed. The dimensionless parameter in Equations 7-10 are taken as *ζω* = 1. and *F*_0_ = 1. Cutoff radius for the GNM is taken as 7.0 Å. The dotted horizontal line shows the equilibrium pair correlation of unperturbed protein given by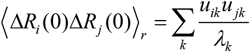.

The upper left panel in Figure 3 compares the time correlation function between Arg55 and 83 when residue 55 is perturbed. The solid curve shows the correlation *C*_55,83_(*ωτ*) between the fluctuation of residues Arg55 and Phe83 where the fluctuations of Phe83 are observed *ωτ* units later than those of Arg55. The lower dot-dashed curve on the other hand shows the correlation *C*_83,55_(*ωτ*). The abscissa covers the 0 ≤ *ωτ* ≤ *π* range. At fixed *ωτ*, fluctuations of residue Arg55 have larger effect on later values of Phe83. The trend is reversed in the next interval *π* ≤ *ωτ* ≤ 2*π* as discussed in the theory section. Throughout this study, we concentrate on the first range only, which represents the slow perturbation range. The *ωτ* = 0 intercept shows the case of static perturbation, which is weaker than the unperturbed correlation between Arg55 and Phe83. As *ωτ* increases, correlation between the pair increases. In the remaining three panels, correlations of residue Arg55 with the other members of the allosteric pathway are shown. In the lower left panel, for the Arg55-Phe113 pair, an increase in static perturbation at *ωτ* = 0 is stronger than the unperturbed correlation. If residue i is perturbed and | *C*_*ij*_(*ωτ*) | > | *C*_*ji*_(*ωτ*)| then the residue i drives j and residue j is driven by i. With this definition, we see from Figure 3 that when Arg55 is perturbed, residues Arg83, Phe113 and Trp121 drive Arg55, but Arg55 drives only Ser99, with a strength that is an order of magnitude weaker.

Next, we analyze the difference *C*_*ij*_(*ωτ*) − *C*_*ji*_(*ωτ*) which shows the directionality of correlations upon perturbation of a residue. For this purpose, we calculated the values of *C*_*ij*_(*ωτ*) for all values of i and j where the residue indicated by the first index is perturbed. Thus, *C*_*ij*_(*ωτ*) is the correlation when i is perturbed and j is observed at a time *ωτ* later, and *C*_*ji*_(*ωτ*) is the correlation when j is perturbed and i is observed *ωτ* later. A positive value of the difference shos that i is the driver residue. Positive values of this difference are shown in Figure 4. Hence, the horizontal axis indicates the driver residue index i

**Figure 4.**
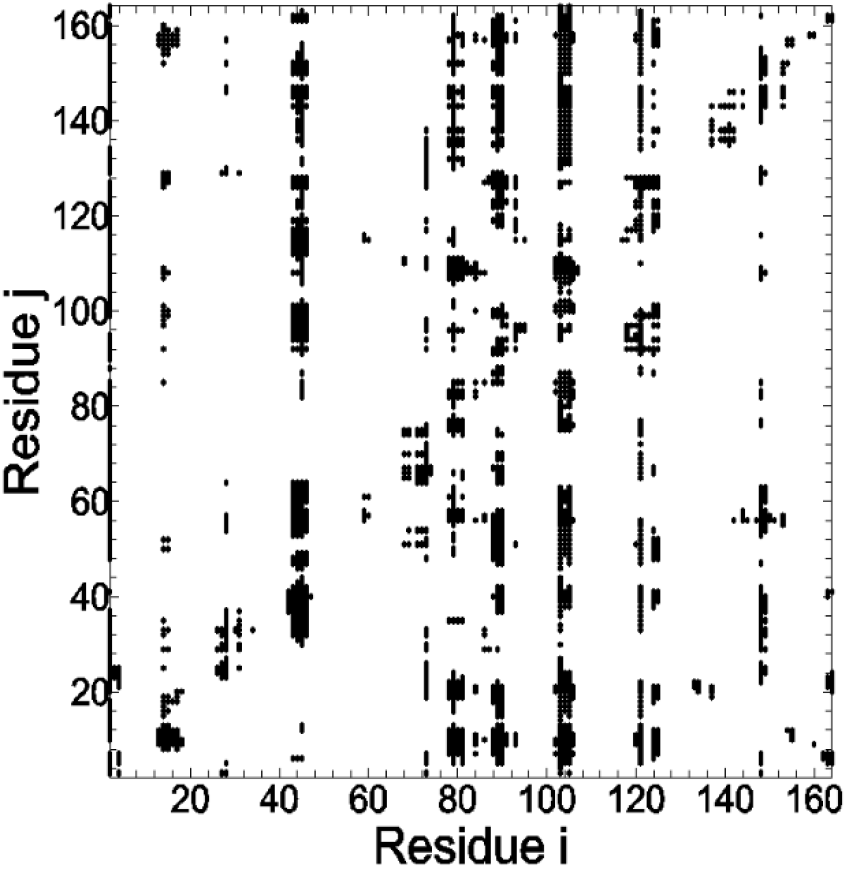
Values of | *C*_*ij*_(*ωτ*) − *C*_*ji*_(*ωτ*)| presented in terms of residue indices i and j. Black dots show values of this difference in the range 0.003 < *C*_*ij*_ − *C*_*ji*_ < 0.07 which covers the top 15% of all points obtained with *F*_0_ = 1, *ωτ* = 0.1, *ζω* = 0.1.

which is perturbed by a cosine force of magnitude *F*_0_ = 1 and *ωτ* = 1.0. The vertical axis indicates the driven residue index j which is affected by the perturbation. The calculations are performed with *ζω* = 0.1. The values of *C*_*ij*_ − *C*_*ji*_ indicate the level of causality between residue pairs. In this example, 0 < *C*_*ij*_ − *C*_*ji*_ < 0.07. There are 13,366 points with positive values. Only the largest 15% of those points are shown in the figure in the interest of identifying important pairs with causality. Interestingly, driver residues are clearly visible as specific vertical patterns in the figure. Accordingly, residues between 42-46, 78-81, 88-91, 100-106, 121, 123-125 and 147-149 act as strong drivers.

Each residue is under the influence of all other residues. This is reflected in the sum 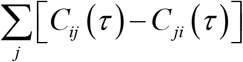. A positive value of this sum indicates that the residue i acts as a driver, and conversely a negative value indicates a driven residue. In Figure 5, the values of this sum are plotted as a function of residue index.

**Figure 5.**
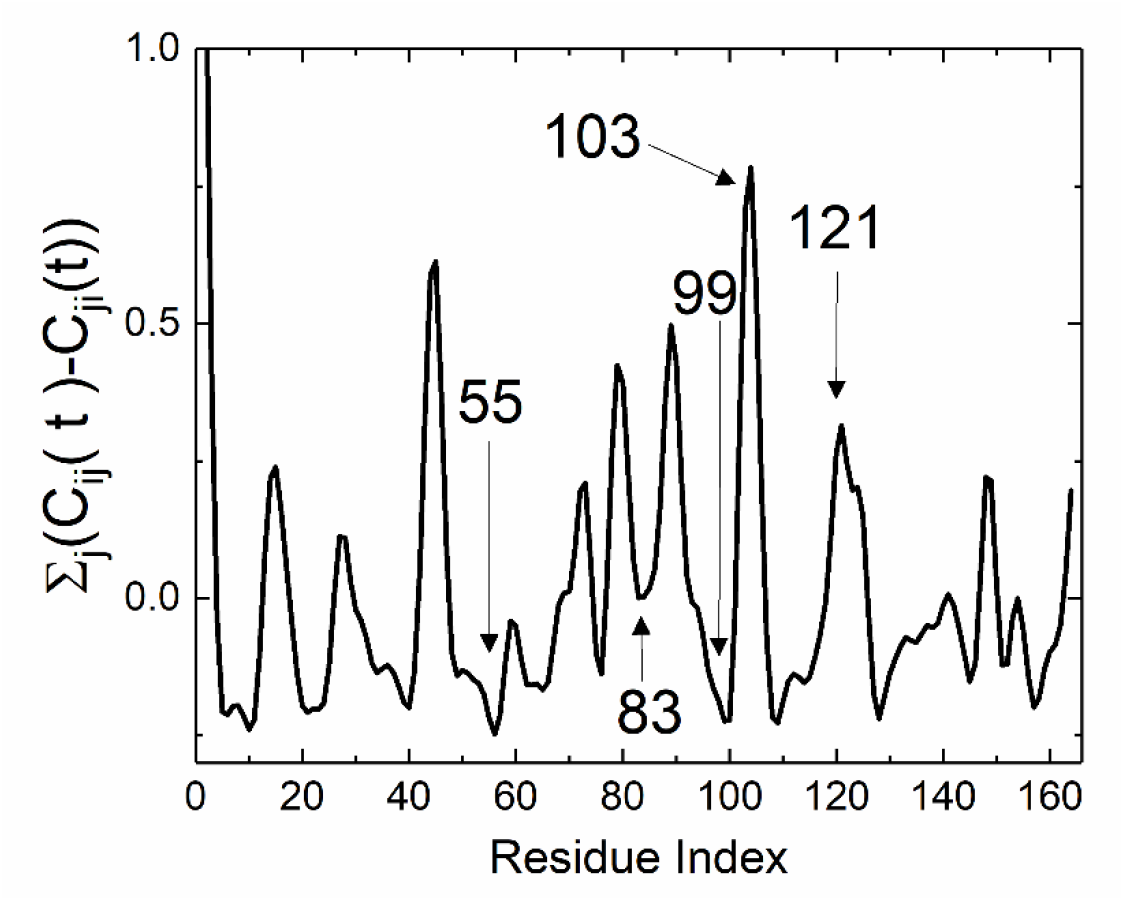
Values of the sum 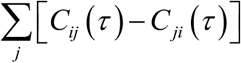 presented as a function of the residue index i. See legend to Figure 4.

The peaks in the figure correspond to strong driver and the minima to driven residues. The deepest minimum is for residue Arg55, which is the active site residue that induces the cis-trans isomerization of the substrate. Among the maxima, residues Glu43, Phe83, Ala101 and Trp121, which are known to play a role in the allostery, are observed.

### 3. Determining the community residues

Understanding how proteins function often boils down to deciphering the intricate networks of interactions between their amino acids. These interactions extend beyond direct physical contacts, forming dynamic communities of residues that act in concert. While the term “community” is widely used, how do we actually identify and understand these hidden groups within a protein? Perturbing a specific residue and observing the other residues which fluctuate with a small phase difference, i.e., with small values of *θ*_*ij*_ is a plausible approach. This synchronized movement suggests a protein community, a cluster of residues that share a functional response. Thus, perturbing a single residue acts as a probe, uncovering the existing communities within the protein. These communities might not be static groups, but rather dynamic ensembles that fluctuate depending on the surrounding environment and cellular needs. However, some communities exhibit remarkable persistence. Despite the location of the perturbation, these core groups consistently show synchronized movements, hinting at their crucial roles in protein function. Identifying these persistent communities through perturbation may help to understand the protein’s functional landscape. In Figure 6, we present the values of *θ*_*ij*_ = *θ*_*j*_ −*θ*_*i*_, as a function of residue index where the number in each figure shows the residue which is perturbed. The calculations are performed with *F*_0_ being the force vector with a 1.0 in the component corresponding to the forced residue and zero otherwise. Below, we used the notation *F*_0_ = 1 for brevity. *ωτ* = 1.0, *ζω* = 1.0.

**Figure 6.**
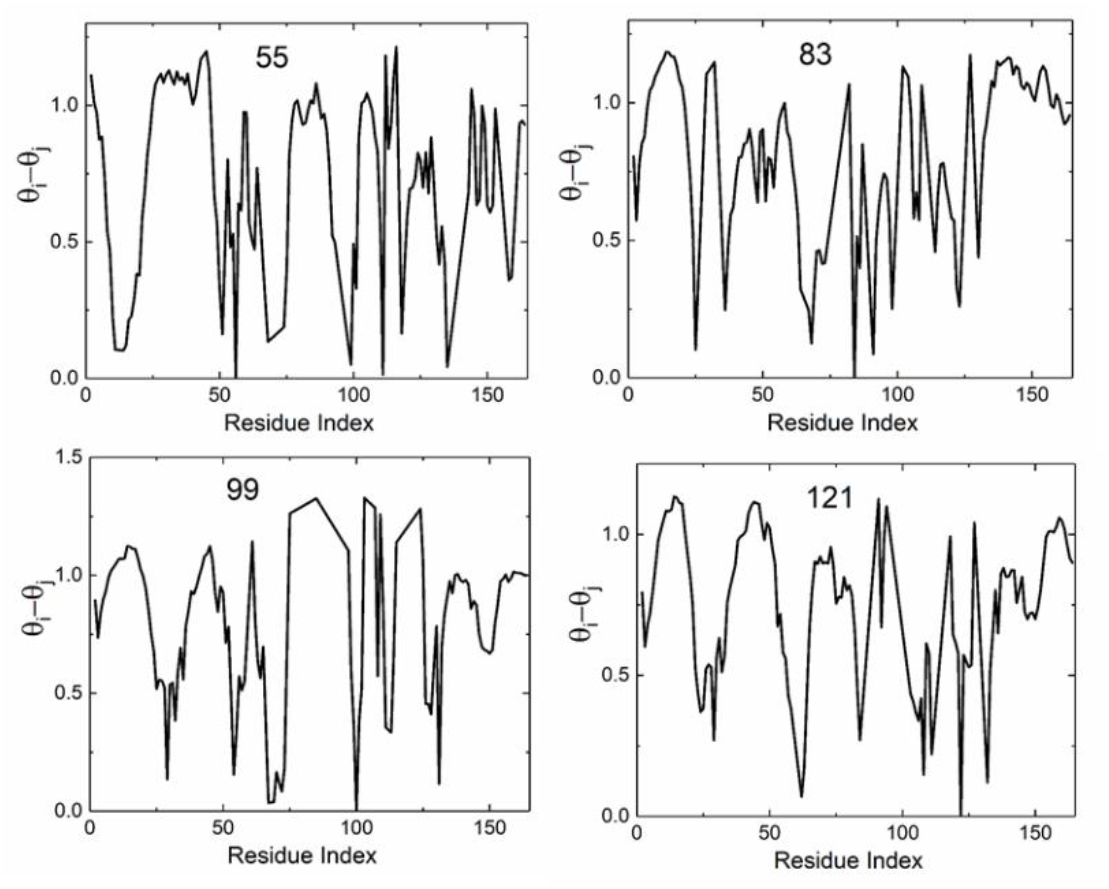
Phase difference between a perturbed residue (identified in each figure) and the remaining residues of the protein, plotted as a function of residue index. Calculations are performed with *F*_0_ = 1, *ωτ* = 1.0, *ζω* = 1.0.

The curves in Figure 6 do not show long stretches of residues for which *θ*_*ij*_ is close to zero, which would indicate same phase with the perturbed residue. However, the curves identify single residues that are in phase with the perturbed residue. For example, in the upper left panel of the Figure, residues Ser99, Phe113 and Val132 move in phase with Arg55. Lower left panel shows that residues Arg55 and Glu74 move in phase with Ser99.

While perturbation with values of *ζω* = 1.0 is unable to identify communities, calculations with a lower frequency perturbation, *ζω* = .01 clearly can identify communities, as may be seen from Figure 7. Several regions of the protein that fluctuate with the same or similar phases are seen. Thus, for all panels the range of residues 30-60 which is the helix-coil-beta motive that contains the catalytic residue Arg55 forms a community commonly observed when residues Arg55, Phe83, Ser99 and Trp121 are perturbed.

**Figure 7.**
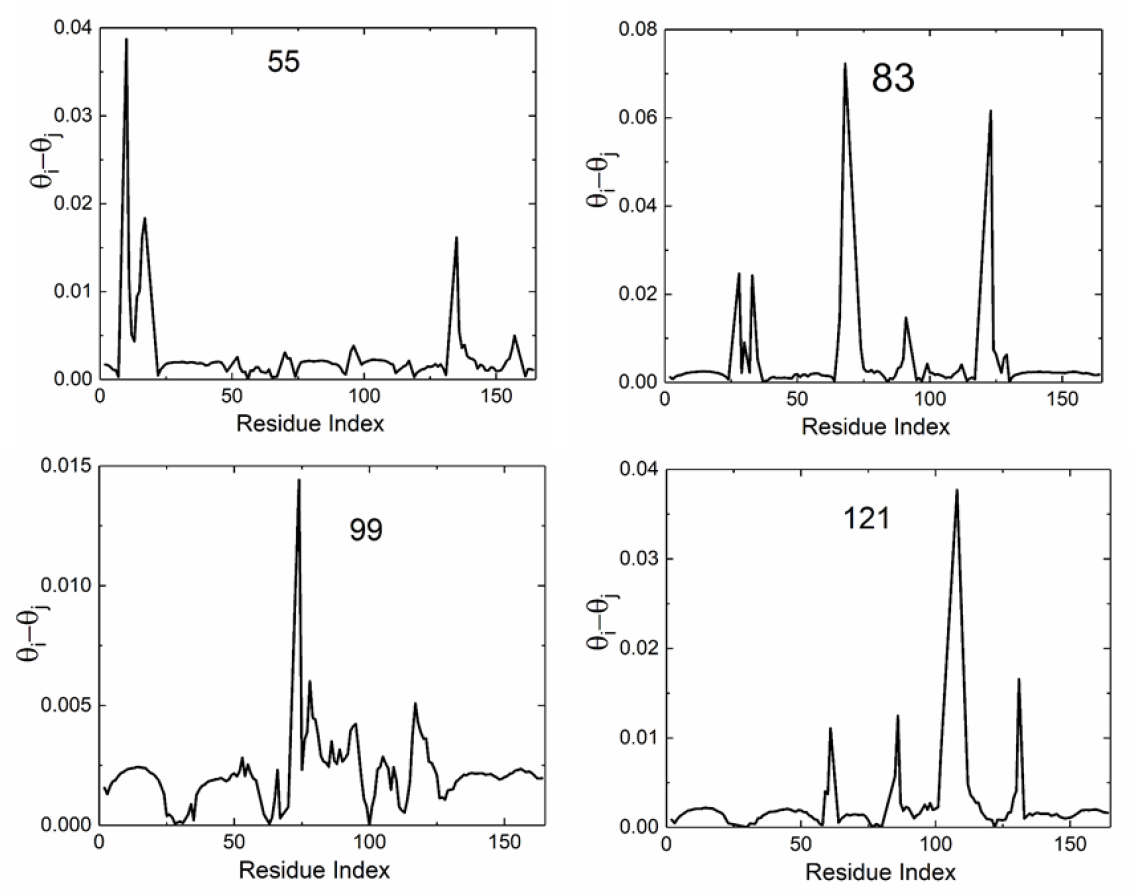
Phase difference between a perturbed residue (identified in each figure) and the remaining residues of the protein, plotted as a function of residue index. Calculations are performed with *F*_0_ = 1, *ωτ* = 1.0, *ζω* = .01.

### 4. Effect of mutation on Cyclophilin activity

In order to see the effects of mutation on the catalytic activity of Cyclophylin A, (i) the distal residue Ser99 was mutated to Thr99 (PDB code 3K0O) and (ii) the central catalytic residue Arg55 was mutated to Arg (PDB code 3K0R). The result was approximately 60 fold decrease of the rate of conversion of cis to trans of the ligand for both structures^28, 29, 39^. In Figure 8 we show the changes in driver-driven relations for the two mutations. A positive value for a residue indicates that mutation makes that residue a stronger driver relative to the wild type. In both panels of Figure 8, we see that Gly80, which is already a driver residue in the wild type structure becomes a stronger driver with either mutation. On the contrary, residues Ala103, Trp121 and Asn149 which are drivers in the wilt type structure, lose this property upon mutation of either Ser99 and Arg55. Overall, it appears that mutation upsets the driver-driven balance in the protein and leads to loss of function.

**Figure 8.**
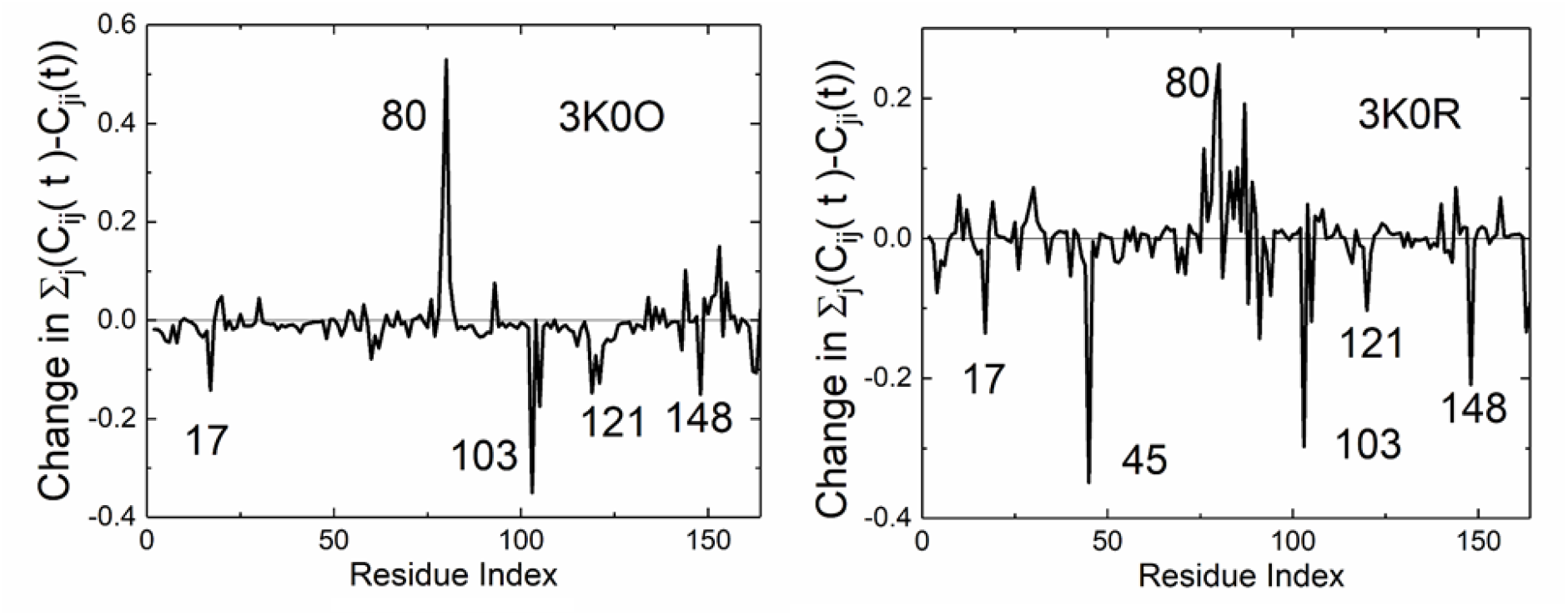
The differences 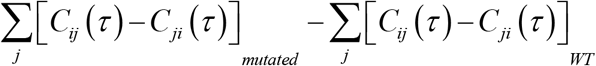 in driver-driven relations for the mutated and wild type structures. The left panel is for the Ser99Thr mutation, the right panel is for the Arg55Lys mutation. Largest changes are indicated in each panel.

## Discussion and Conclusion

The application of DP-GNM to Cyclophilin A has facilitated a comprehensive analysis of its dynamic behavior and provided insightful details into the temporally ordered events within its allosteric network. Our results reveal a complex interplay between driver and driven residues, organized within a hierarchical framework of information flow. Key residues such as Arg55 and Trp121 exhibit a strong driver character, exerting significant influence on others, while some, like Phe113, demonstrate a dynamic role, transitioning between driver and driven depending on the context. This suggests a cascading pattern of information transfer within the protein, where perturbations at specific sites trigger a sequential propagation of conformational changes throughout the network.

Intriguingly, residue 55, traditionally viewed as the primary driver of allosteric regulation due to its active site role, primarily appears as a driven residue in our analysis. This suggests that its functionality is largely governed by conformational changes initiated elsewhere in the protein, highlighting the intricate web of influence between distinct regions within Cyclophilin A. This finding underscores the limitations of the single-driver model of allosteric regulation and emphasizes the need to account for the complexity of hierarchical information flow within proteins.

Furthermore, DP-GNM has identified the presence of dynamically coupled communities—groups of residues exhibiting synchronized movements upon perturbation. These communities are likely to represent functionally relevant modules within Cyclophilin A, responsible for specific aspects of its activity. Identifying these communities enhances our understanding of protein function and the underlying dynamic pathways. However, it is crucial to acknowledge that these communities are not static entities. They may reconfigure under different conditions such as frequency of perturbation or upon binding ligands, highlighting the dynamic nature of protein structure and function. Further research is necessary to explore the plasticity of these communities and their relationship to protein function under various environments.

While directionality exhibits a periodic reversal within intervals of (n-1)π < wt < nπ, it’s important to note that the first interval, 0 < wt < π, holds particular relevance for understanding slow dynamics. If perturbations primarily reside within this range, the observed directionality remains consistent, and concerns regarding potential reversals within subsequent intervals become less pertinent.

Limitations of the current study include the use of a simplified GNM model that assumes isotropic fluctuations and neglects specific intramolecular interactions. Additionally, the chosen frequency range focuses on slower processes and might not capture the full spectrum of allosteric dynamics. Future work can incorporate anisotropic models and explore a wider range of frequencies to achieve a more comprehensive understanding of protein dynamics and the complete repertoire of information flow within proteins.

DP-GNM has emerged as a powerful tool for elucidating the intricacies of protein dynamics and allosteric regulation. By revealing the temporal sequence of events, directionality of information flow, and dynamic interplay between different regions, DP-GNM provides valuable insights into the mechanisms underlying protein function. Our analysis of Cyclophilin A demonstrates the hierarchical organization of the allosteric network, the driven nature of the active site residue Arg55, the existence of dynamically coupled communities, and the importance of considering perturbation frequency ranges for interpreting directionality. This approach has the potential to advance our understanding of protein function and provide a foundation for the development of novel therapeutic strategies targeting allosteric processes. Future research directed towards refining the model, exploring a wider range of proteins, and integrating experimental data will further solidify DP-GNM as a useful tool in the field of protein dynamics and open new avenues for understanding and manipulating protein function.

While this study focused on DP-GNM’s application to a single case of catalytic activity of Cyclophilin A, its potential extends to analyzing systems under various perturbations, including ligand binding, for example. A ligand docked at a specific site on the protein with its dominant frequency overlapping with the protein’s vibrational modes, could be efficiently absorbed, effectively “pumping” energy into the protein’s internal network. This vibrational energy transfer could act as a periodic perturbation, triggering a ripple effect through the protein structure. DP-GNM could identify the key residues that initially “catch” the vibrational energy from the ligand, the communities of residues that synchronize their movements in response, and ultimately, the downstream functional consequences of this allosteric excitation. This detailed understanding of ligand-induced dynamics could offer valuable insights into ligand efficacy, selectivity, and even potential off-target effects. Moreover, by analyzing the directionality of information flow under ligand binding, DP-GNM could inform the design of ligands with targeted vibrational properties. By precisely engineering ligand vibrations to resonate with specific protein modes, we could potentially steer allosteric responses towards desired outcomes, paving the way for novel strategies in drug design and protein engineering.

Intriguingly, the shifts in driver/driven residues, emergence of synchronized communities, and coordinated responses under dynamic loading bear resemblance to the dynamic nature of the circadian clock. While the circadian clock primarily relies on biochemical processes for its rhythm generation, several types of mechanical perturbations can influence its function in indirect ways. This raises the question: could DP-GNM, applied to protein networks during circadian oscillations, illuminate the interplay between environmental cues and internal protein dynamics?

## Supplementary Material

We consider the system

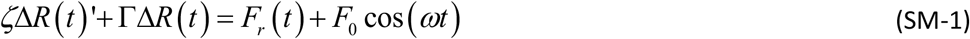

where *F*_*r*_(*t*) is a random force and *F*_0_ is a constant vector. *ζ* is the friction coefficient, and Γ is the spring matrix of the Gaussian Network Model, GNM^3^. *F*_*r*_(*t*) is a zero mean white noise satisfying 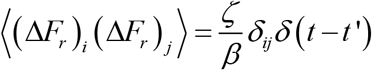 where, *δ* is the Kronecker delta, *δ*(*t* − *t*’) is the Dirac Delta function and *β* = 1/ *k*_*B*_*T*0, *k*_*B*_ being the Boltzmann constant and *T*_0_ the absolute temperature.

Solutions of (SM-1) are of the form

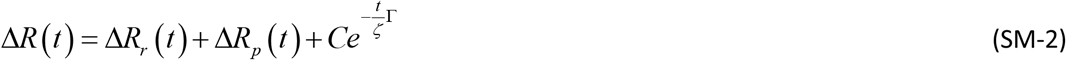

with Δ*R*_*r*_(*t*) and Δ*R*_*p*_(*t*) being solutions corresponding to the random force *F*_*r*_(*t*) and deterministic forces, respectively. The term 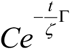 with an arbitrary constant vector *C* corresponds to the solution of the homogeneous equation, i.e., with zero force. As we are interested in the correlations

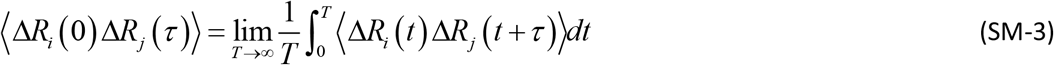

the term 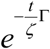 will not contribute as it decays rapidly as *t* →∞, hence we will neglect it.

To evaluate Δ*R*_*r*_, we consider the Fourier series expansion for *F*_*r*_ in the interval [0; 2T]

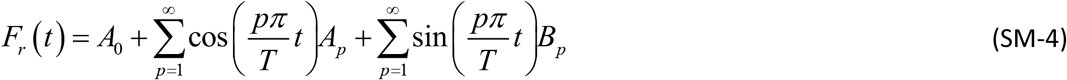

with the coefficients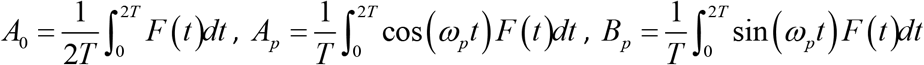

The random solution Δ*R*_*r*_(*t*) then turns out to be (see 2004 for further details)^4, 5^

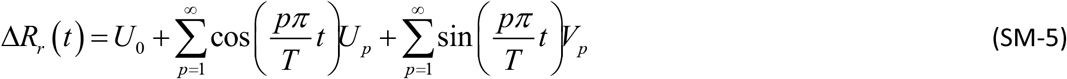

where,

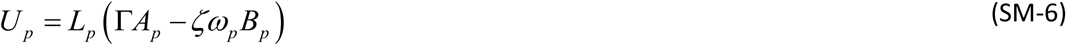

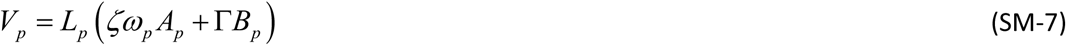

where 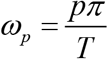 and

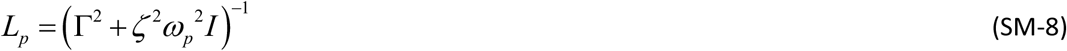

Since *F*_*r*_(*t*) has zero mean, it follows that the coefficients *A*_*p*_, *B*_*p*_, *U*_*p*_, *V*_*p*_ and thus Δ*R*_*r*_(*t*) are all zero mean random processes. To compute the correlation, we first simplify the terms

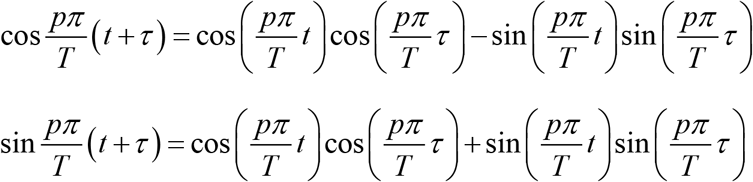

to get

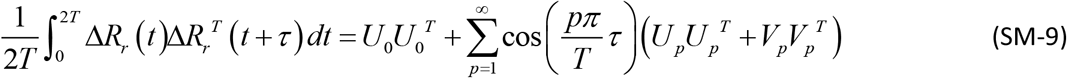

We take ensemble averages

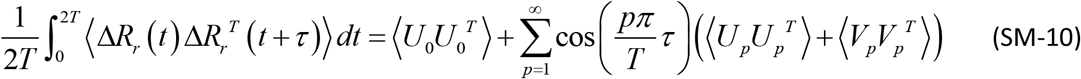

We then have

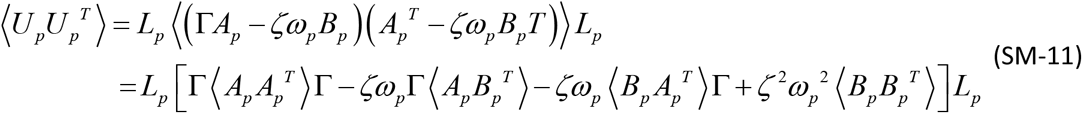

We now compute the ensemble averages 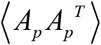, etc.

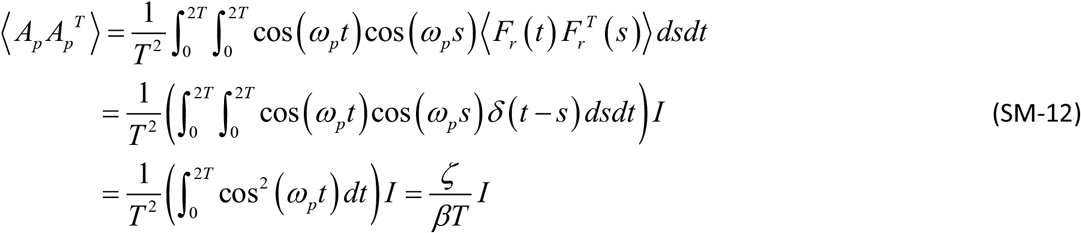

with the identity matrix *I*. Similarly,

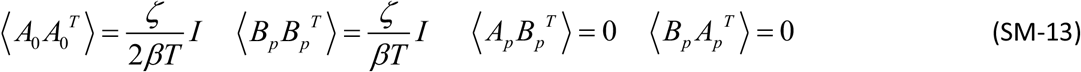

This in turn gives

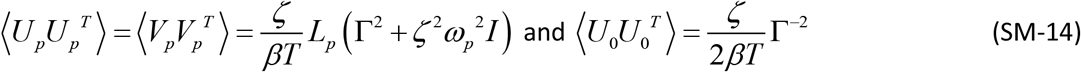

where,

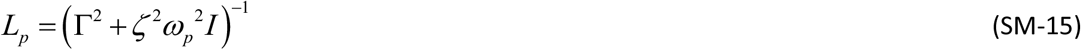

The second equality follows similarly. Then

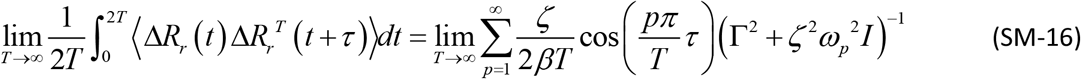

Noting that 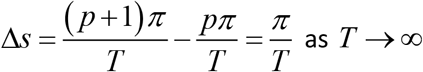 the sum converges to the integral on the interval (0, 2*T*) and then on (0, ∞). Summing up we get

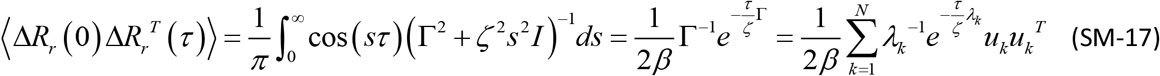

We now look at the periodic, deterministic term. Similar to the computations above, we get

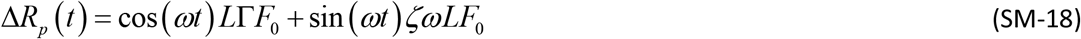

With

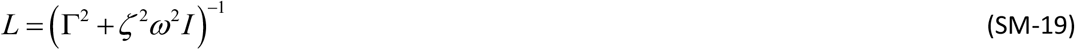

It will be more informative to look at the components

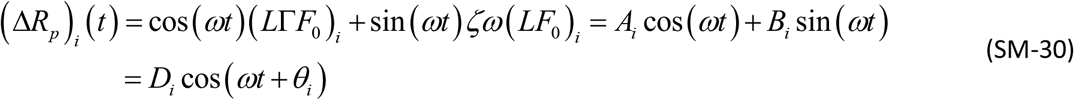

with 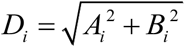 and 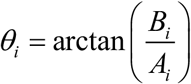. The correlation integrals then can be computed to yield

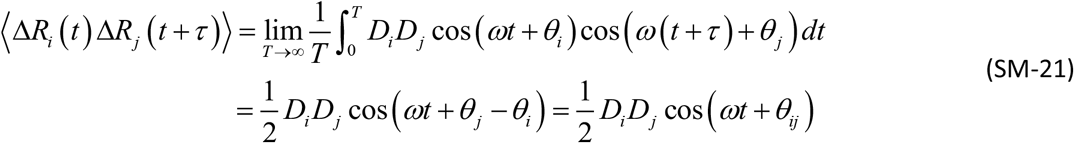

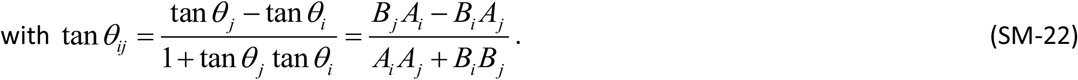

Finally, as Δ*R*_*r*_(*t*) is a random process with zero mean, all the mixed terms in the correlation will vanish to give:

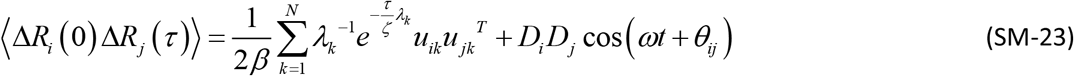

